# *Drosophila* Hox genes induce melanised pseudo-tumours when misexpressed in hemocytes

**DOI:** 10.1101/363978

**Authors:** Titus Ponrathnam, Rakesh K Mishra

## Abstract

Homeotic genes are the key early determinants of cell identity along the anterior-posterior body axis across bilaterians. More recently, however, several late non-homeotic functions of hox genes have emerged in a variety of organogenesis processes, including in mammals. Being crucial factors in determining cell identity and organogenesis, the misregulation of hox genes is likely to be associated with defects in these processes. Several studies have reported misexpression of hox genes in a variety of malignancies including acute myeloid leukaemia. Considering that *Drosophila* is a well-established model for the study of haematopoiesis, we ectopically expressed the hox genes, *Dfd*, *Ubx*, *abd*-*A* and *Abd*-*B*, to ask if and how it will alter the process of haematopoiesis. We observed black melanised spots circulating in the viscera of the larvae and extensive lethality at during the pupal stage in these conditions. Such abnormalities are the hallmark of dysregulated haematopoiesis. We also observed an increase in blood cell number as well as their enhanced differentiation into lamellocytes. Our study opens a new possibility of addressing the function hox genes in normal and leukemogenic hematopoiesis with potential implications in downstream targets for diagnostic markers and therapy.

**Summary:** *Drosophila* Hox genes, when expressed in blood cells, are leukemogenic, induce cell autonomous proliferation and differentiation. This reinforces previous studies in vertebrates and allows for Hox induced leukaemia to be studied in *Drosophila.*

## Introduction

One of the most striking aspects of life is the variety in body forms. Despite this variety, there is an even more striking similarity at the genetic and molecular level, in the developmental mechanisms that lead to variety across the species. For example, in spite of the evolutionary distance of hundreds of millions of years between vertebrates and *Drosophila*, many organ and tissue types show a degree of homology with each other and much of the key developmental pathways are highly conserved.

The hematopoietic system is no exception to this conservation. Hemocytes of *Drosophila* resemble the myeloid lineage of blood cells [1]. The most abundant cells, plasmatocytes, are the equivalent of macrophages that are involved in a variety of processes, such as responses to pathogens, removal of apoptotic cells, deposition of the extracellular matrix during embryonic development, etc. [2]. The next most abundant cells are the Crystal cells, specialised to induce melanisation reactions in the presence of pathogens and wound healing [3], resemble the granulocytes, and contribute about four per cent of the blood cells. Lamellocytes are the least abundant population of blood cells, usually only appearing in circulation upon the larva being challenged by any object too large to be cleared off by the macrophages, for example, parasitoid wasp eggs [4,5].

The extent of this resemblance includes the molecular pathways involved in the development of these cell types. *Serpent*, [6] is related to GATA 1, 2 and 3 of vertebrates. GATA-2 is responsible for blood progenitor proliferation and survival [7,8]. GATA-1 is required for progenitor differentiation into erythrocytes, megakaryocytes and eosinophils [9–11]. Similar to GATA-2, *Srp* is required for progenitor maintenance and proliferation. Loss of function in *Srp* leads to a reduced number of progenitors and a loss of all hemocytes. It is also required in plasmatocyte differentiation, similar to GATA-1 [12]. Additionally, *ushaped* is related to the Friend of GATA (FOG) family. FOG-1 and GATA-1 are required together for erythrocyte and megakaryocyte differentiation [13,14]. FOG-1 interacts with GATA-1 to represses eosinophil differentiation and must be downregulated for eosinophil differentiation [15]. Similarly, *ush* is expressed in hemocyte precursors and plasmatocytes, and must be downregulated for crystal cell development [16].

Various signalling pathways involved in regulating hematopoiesis, as would be anticipated, are conserved between vertebrates and *Drosophila.* Examples include Jagged-1, the vertebrate homolog of *Serrate*, a ligand of Notch, is produced by the stromal cells of the bone marrow, to regulate HSC proliferation and survival [17]. *Ser* performs a similar role, being secreted by cells of the Posterior Signalling Centre [18], a set of regulatory cells at the posterior end of the Lymph Gland (LG). Vertebrate JAK2 is required for erythropoiesis [19], while STAT5 is required for proper progenitor and myeloid cell function [20]. The *Drosophila* JAK/STAT pathway is required within the LG for the maintenance of prohemocytes, among other things [21]. Transformations in JAK2 can lead to leukemogenesis in vertebrates [22], similar to how gain of function *hop* mutants behave [23]. The Toll pathway is also conserved, playing a major role in innate immunity in both vertebrates and flies [24].

One aspect of vertebrate hematopoiesis that has not been mirrored in *Drosophila* is the role of Hox genes. Hox genes are well known for their conserved role in body axis formation across all bilaterians[25], but also play roles in vertebrate hematopoiesis [26], autophagy [27], as well as cell proliferation, differentiation, migration and apoptosis [28]. Hox genes are transcribed in HSCs as well as lineage progenitors, while being suppressed in differentiated blood cells [29–33]. Overexpression models lead to blockages in certain stages of development, expansion of HSCs, the circulation of blast cells, etc. [34–39]. On the other hand, in *Drosophila*, other than *Antp*, which is implicated in setting up the location of the LG [40], as well as later marking the PSC [41] hox genes are not known to play any major role.

In this study, we show that overexpression of hox genes, *Dfd*, *Ubx*, *abd*-*A* and *Abd*-*B*, in the blood cells not only leads to melanised bodies, but also to a significant increase in blood cell number and the induction of lamellocyte differentiation. These findings have implications in understating the biological events associated with leukaemia in humans which may open new possibilities of markers and therapy.

## Material and methods

### Fly culturing

Flies were cultured in standard cornmeal and sucrose agar. The wild-type flies used in this study were Canton-S. Flies were maintained at 25°C. The Gal4 driver lines used in this study were obtained from the Bloomington stock centre. The UAS lines used in this study, viz., *UAS*-*Dfd*-*HA*, *UAS*-*Ubx*-*HA*, *UAS*-*abd*-*A*-*HA* and *UAS*-*Abd*-*B*-*HA* were sourced from Yacine Graba’s lab. Flies were allowed to lay eggs for 6 hours before being transferred to a fresh vial. Larvae were sacrificed at 96-102 hours post egg lay, before the onset of metamorphosis.

### Immunostaining

For staining proliferative cells we made use of Anti-PhosphoHistone 3 at serine 10, from Upstate (cat# 07-212, 1ng/μL). For confirming the presence of lamellocytes, we used anti-*myospheroid* (DSHB #CF.6G11, 27pg/μL). Blood cells were prepared using an established protocol [42]. Blood cells numbers were quantified using a modified version of the protocol by Petraki, Alexander, & Bruckner, 2015 [43].

## Results

When we expressed the UAS lines different hox genes, *Dfd*, *Ubx*, *abd*-*A* and *Abd*-*B*, under two hemocytes specific drivers, *He*-Gal4 and *Hml*-Gal4, we observed melanised spots. Such melanised spots are absent when a driver specific to the fatbody, *Lsp2*-Gal4, is used but found when a driver expressing in both the fatbody and hemocytes, *cg*-Gal4, is used. We also observed the degree of penetrance is in proportion to the strength of the respective drivers in hemocytes (Figure 1). Taken together, this implied that the melanised pseudo-tumour phenotype we observe is of hemocyte origin.

**Fig.1.**
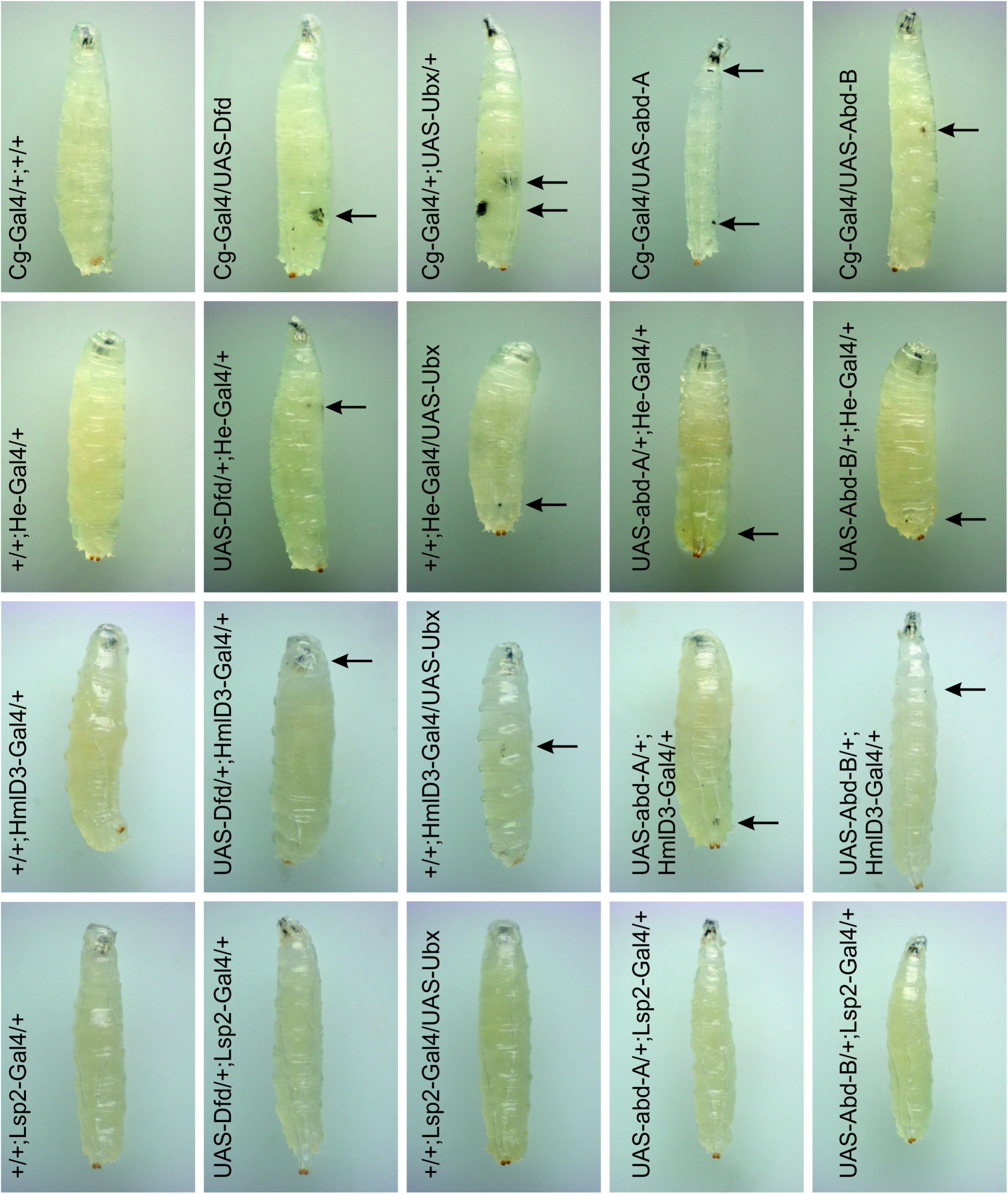
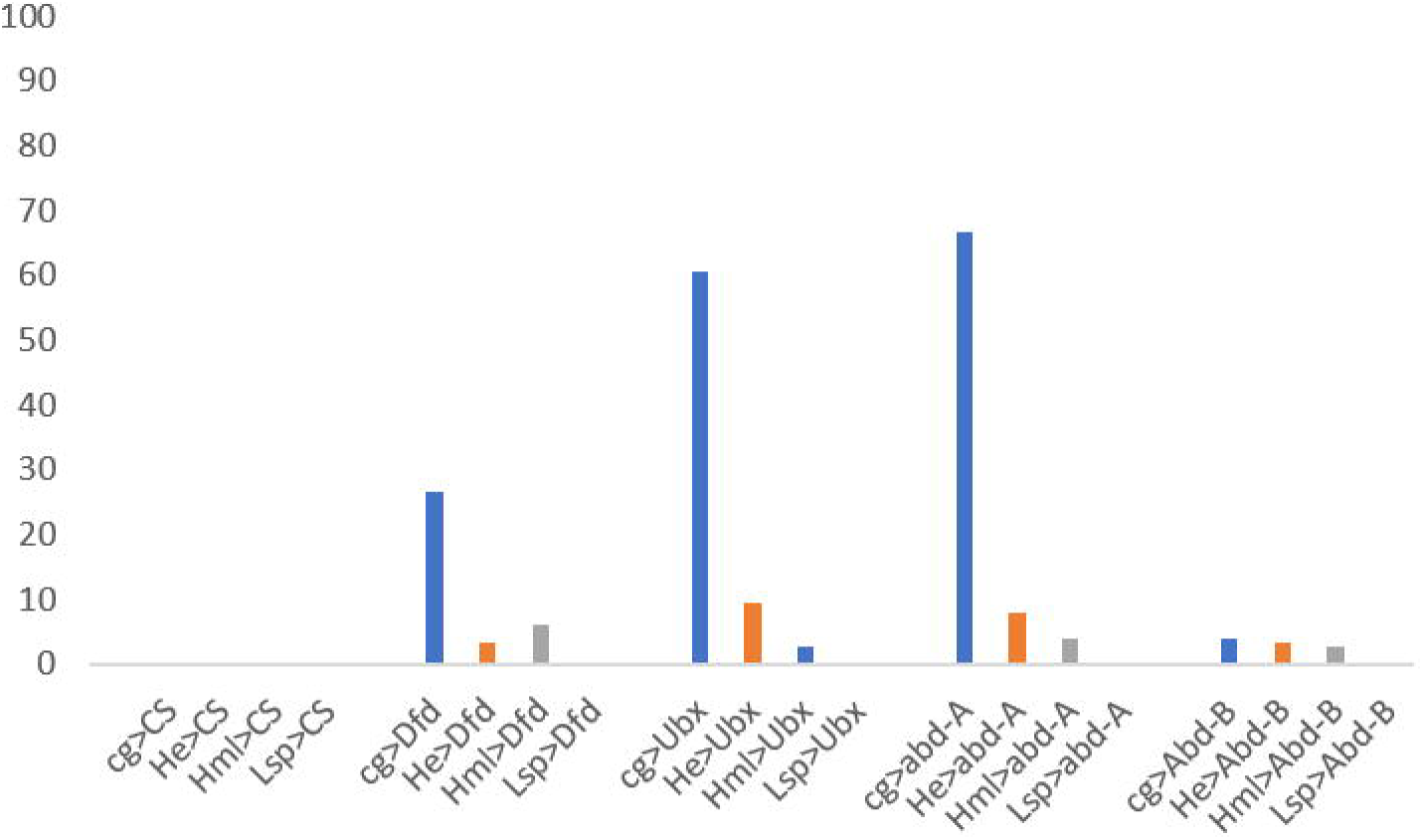
A) Larvae with subcutaneous tumors. *Dfd*, *Ubx*, *abd*-*A* and *Abd*-*B*, when expressed under the drivers *cg*, *He* and *Hml* lead to melanised bodies in the viscera. B) The size and penetrance of these bodies was maximum when expressed under *cg*. While tumors do manifest when *He* and *Hml* are used, they are much rarer and smaller. Expression under *Lsp2*-Gal4 does not lead to the formation of such bodies.

We quantified the number of blood cells in our overexpression lines using a modified version of established methods [42,43]. Larvae were dissected in fixed volumes of PBS in 8mm chambers; blood cells were allowed to settle, fixed and then visualised with phalloidin and made a relative estimation of the number of blood cells in each larva by imaging five fields of view and summing up the number (Figure 2). Our results show that when under the expression of the fatbody driver *Lsp2*, the number of hemocytes was comparable to the controls. Blood specific expression, however, led to a significant increase in the number of circulating hemocytes. Interestingly, while the penetrance of melanised spots was lower, blood specific drivers showed a larger number of blood cells (Figure 2).

**Fig.2.**
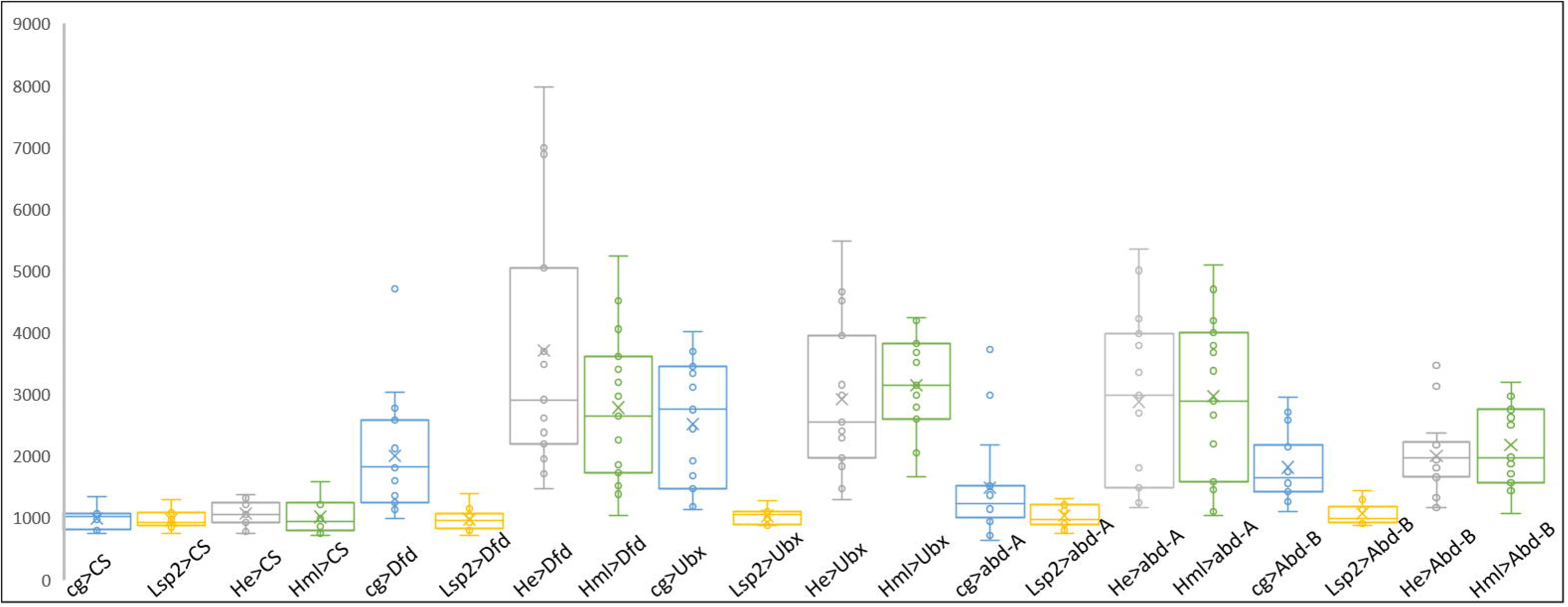
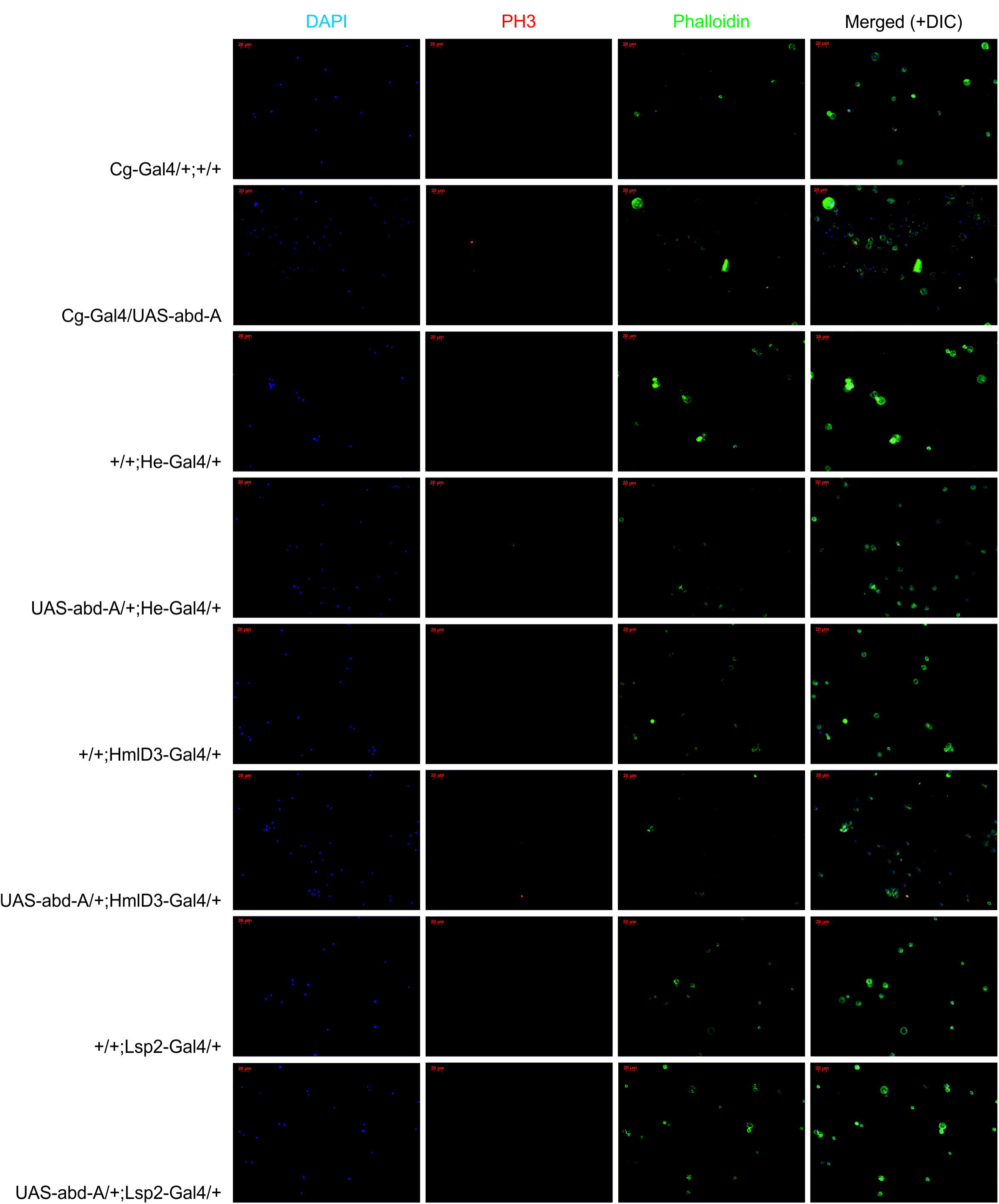
A) Quantification of blood cells. When Dfd, Ubx, abd-A and Abd-B are driven in blood cells under *cg*-Gal4, *He*-Gal4 or *Hml*-Gal4, but not in the fatbody (Lsp2-Gal4) there is an increase in the overall cell number. B) PhosphoHistone3+ nuclei appear when these genes are expressed in the blood cells, indicating that some of the increase in cell number maybe due to cell autonomous proliferation.

Previous studies have shown that cells of the LG do not enter into circulation until the onset of metamorphosis. However, *Hml* and *cg* express in the cortical region of the LG, and *He* expresses throughout. Thus, the question arose as to whether the increase in cell number was due to an increase in cell proliferation at the LG or were circulating cells proliferating in a cell-autonomous manner. To this end, we checked for the presence of the mitotic cell marker PH3. Again, we observed cells positive for PH3, when Hox genes were expressed in the blood cells, and not when expressed exclusively in the fatbody (Figure 2, Supplementary Figure 1). Unlike previous reports, we did not find proliferative cells in our controls [44]. This may be due to a loss of cells in our preparations or more robust immunostaining on our part. Thus, while we cannot rule out the possibility that LG cells contribute to this increase, at least a fraction of the increase takes place due to the cell autonomous division of Hox overexpressing cells.

While imaging the blood cells, we noticed that there were larger, flattened cells in circulation, reminiscent of lamellocytes. We wished to know if these were bona fide lamellocytes, to which end we stained blood cells for the lamellocyte marker *myospheroid* (Figure 3, Supplementary Figure 2). In control larvae, there were none, or if present, a handful of lamellocytes in circulation. In our overexpression lines, however, we noticed that not only were lamellocytes *mys*+, but so were some of the plasmatocyte like cells. None of the plasmatocytes in the control flies or those overexpressing *Abd*-*B* showed *mys.* Thus, we can speculate that these cells, upon Hox overexpression, are pushed toward the lamellocyte fate.

**Fig.3.**
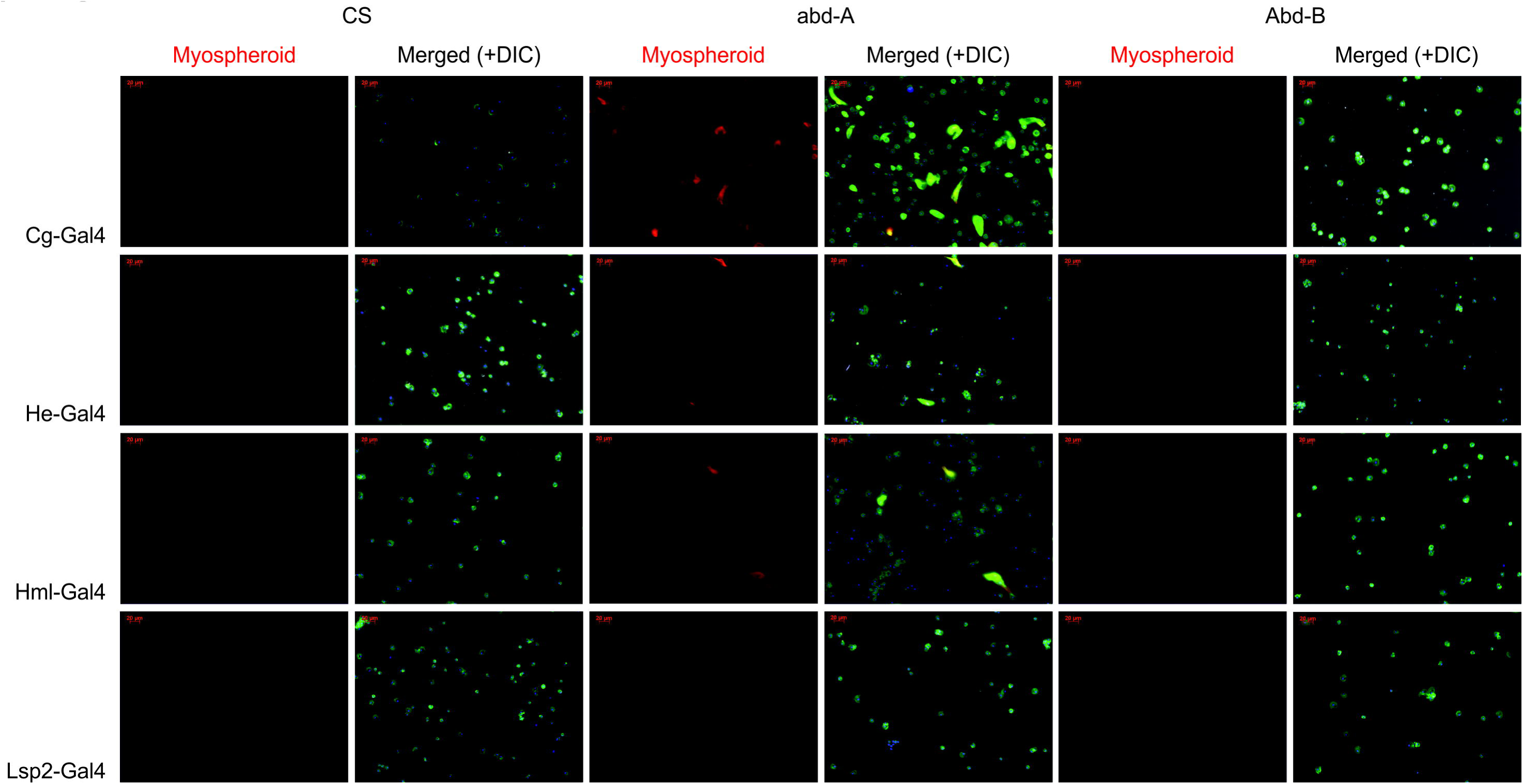
*Myospheroid* staining for lamellocytes. When *Dfd*, *Ubx*, *abd*-*A* but not *Abd*-*B* are driven in blood cells (under *cg*-Gal4, *He*-Gal4 or *Hml*-Gal4), but not in the fatbody (*Lsp2*-Gal4), large, dorsoventrally flattened cells begin to appear in circulation. These stain positive for *mys.* Some circulating plasmatocytes also appear to *mys*+. This indicates that they might be in the process of differentiating.

We also noticed that many pupae were not eclosing when the *cg* driver was used. It has previously shown that aberrant blood cells can induce pupal lethality [45]. However, while we did observe some pupal lethality when the Hox genes were expressed under *He* and *Hml*, the greatest recapitulation of this lethality was when the *Lsp2* Gal4 driver was used (Figure 4). This may be on account of a previous result that suggests Hox genes are repressors of autophagy [27].

**Fig.4.**
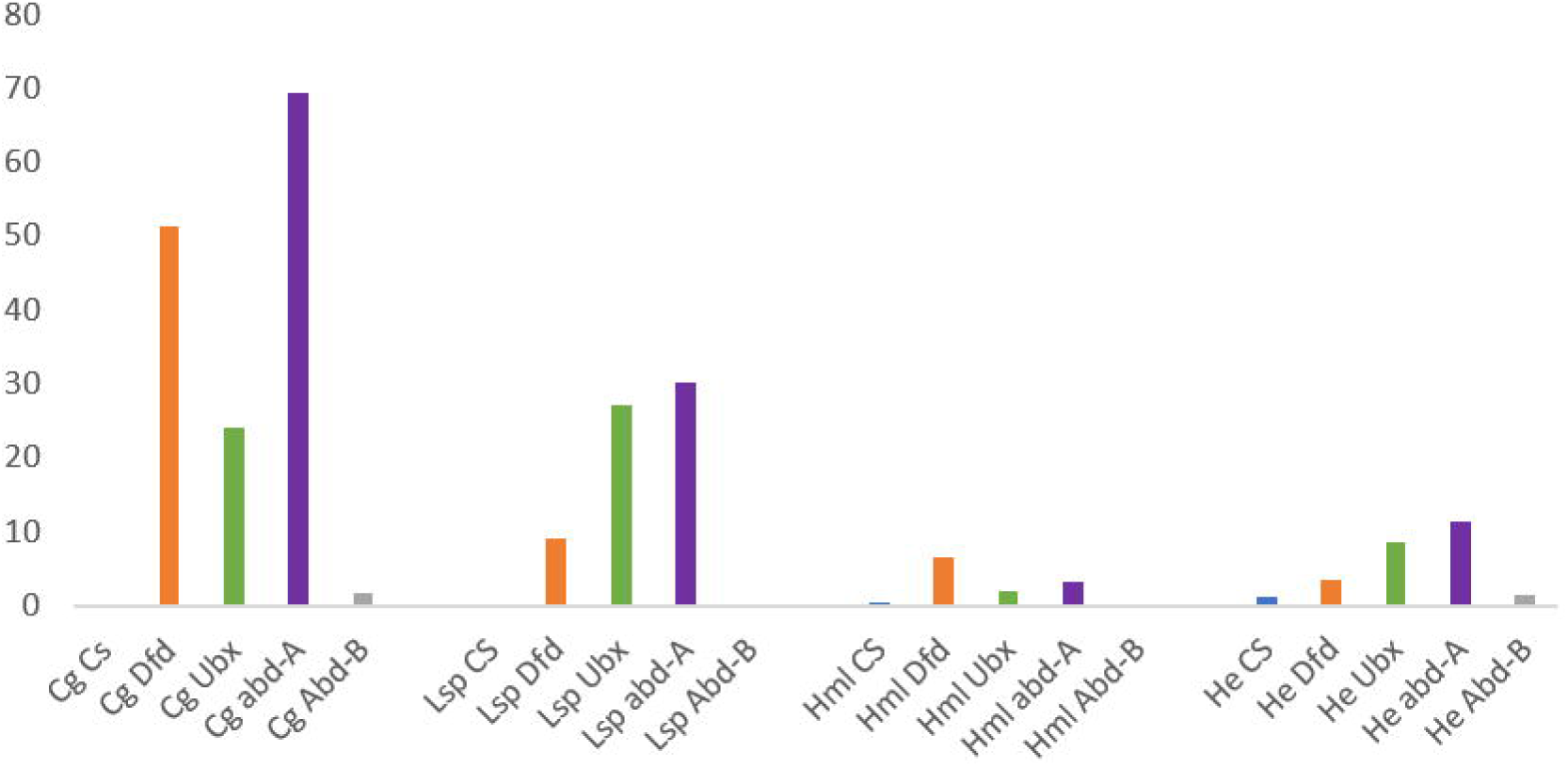
Percentage of pupal lethality, indicated by larvae that fail to eclose. While diving the genes *Dfd*, *Ubx*, *abd*-*A* and *Abd*-*B* with *cg*-Gal4 does cause lethality, so does expressing them in the fatbody under *Lsp2*-Gal4. Driving these genes in the blood cells (*He*-Gal4 and *Hml*-Gal4) leads to a much lower penetrance of this phenotype.

## Discussion

We show that the four Hox genes used in this study in *Drosophila* are capable of inducing melanised bodies in circulation when expressed in the blood cells. These melanotic spots appear only when expressed with Gal4 driver lines active in the LG and circulating blood cells. These spots, therefore, are likely to be aggregations of hemocytes mounting an ectopic immune response. The expression of the genes also triggers cell proliferation. The cells appear to divide in a cell-autonomous manner, which is reflected in the detection of PH3+ cells in circulation. The presence of *myospheroid* positive elongated cells, seen in circulation, also suggests that Hox overexpression leads to the differentiation of the circulating blood cells into lamellocytes. While overexpression of *abd*-*A* shows relatively stronger phenotypes described here, *Abd*-*B* overexpression does not. It supports our earlier finding in which we observed a non-homeotic growth promoter role of *abd*-*A* during formation of the adult cuticle during pupation, while *Abd*-*B* does not show any such role [46].

Our results indicate that Hox genes are causal in leukaemia, reinforcing previous studies in vertebrate model systems, and extending these findings to *Drosophila.* This also opens the possibility that Hox gene induced leukaemias, especially those of the myeloid lineage, can be studied and modelled in *Drosophila.* Till date, the only known Hox gene to participate in hematopoiesis is *Antp*, which marks in the PSC, and provides spatial signals for the development of the LG. In vertebrates, Hox genes have been shown to express within progenitor cells and are rapidly switched off during cell maturation. As our overexpression lines perturb both cell number and differentiation, it is likely that multiple *Drosophila* Hox genes are involved in finetuning the precise programme of *Drosophila* blood cell development.

The fact that the cells appear to be phenotypically confined to plasmatocytes and lamellocytes implies that expression of these genes works in tandem with, and above the specific programme of the cell types. It would be interesting to know which genes are being modulated in our overexpression lines, by profiling their transcription states as well as the binding sites of the individual Hox proteins. In the absence of this information, we speculate that Hox gene overexpression leads to the aberrant transcription of genes. It is known from previous studies that Hox dysregulation in leukaemia is usually concomitant with gain or loss of function mutations in upstream regulators, most commonly Mixed Lineage Leukemia-1 (MLL-1) fusion proteins [47,48], or loss of function EZH2 mutations [49]. It has been reported that MLL-1 fusion proteins have the lowest number of co-operating mutations to induce leukemogenesis [50]. Taken together, we speculate that these driver mutations induce Hox gene activation, which in turn induce leukaemia via aberrant transcription.

## Acknowledgements

We acknowledge Yacine Graba for the UAS lines used, Ravindra Chakravarthi, C Subbalakshmi, Aprotim Majumder and Kasavan S for access to and help with imaging facilities.

